# Antithymocyte globulin inhibits CD8^+^ T cell effector functions via the paracrine induction of PDL-1 on monocytes

**DOI:** 10.1101/2022.07.26.501584

**Authors:** Dragan Copic, Martin Direder, Katharina Klas, Daniel Bormann, Maria Laggner, Hendrik Jan Ankersmit, Michael Mildner

## Abstract

Antithymocyte globulins (ATG) are T cell depleting antibodies used in solid organ transplantation for induction therapy in sensitized patients with high risk of graft rejection. Previously described effects besides depletion of T cells suggest additional modes of action and identified further cellular targets. Here, we examined the transcriptional changes arising in immune cells from human blood after ex vivo stimulation with ATG on a single cell level to uncover additional mechanisms by which ATG regulates T cell activity and effector functions. Analysis of the paracrine factors present in plasma of ATG-treated whole blood revealed high levels of chemokines and cytokines including Interferon-γ (IFN-γ). Furthermore, we identify an increase of surface expression of programmed cell death 1 ligand 1 (PDL-1) on monocytes mediated by the released paracrine factors. In addition, we show that this induction is dependent on activation of JAK/STAT signaling via binding of IFN-γ to Interferon-γ receptor 1 (IFN-γR1). Lastly, we demonstrate that the modulation of the immune-regulatory axis of Programmed cell death protein 1 (PD1) on activated CD8^+^ T cells with PDL-1 found on monocytes mediated by ATG potently inhibits effector functions including proliferation and granzyme B release of activated T cells. Together our findings represent a novel mode of action by which ATG exerts its immunosuppressive effects.

**One Sentence Summary:** ATG increases PDL-1 on CD14^+^-monocytes and inhibits T cell effector functions.

## INTRODUCTION

Induction and maintenance of immunosuppressive therapy have strongly contributed to reduce graft rejections after solid organ transplantation (*1*). In patients with high immunological risk of acute graft rejection antithymocyte globulin (ATG) is recommended for induction therapy (*2, 3*). ATG is a mixture of purified polyclonal γ-globulins obtained from sera of horses or rabbits inoculated with cells from human thymus (*4*). These antibodies bind to a vast array of antigens found on the cell surface of human T lymphocytes as well as other immune cells (*5*). ATG depletes T lymphocytes via classical complement-dependent cell cytotoxicity (CDCC), antibody-dependent cell cytotoxicity (ADCC), opsonization of reactive T cells and the induction of the activation-induced cell death pathway (AICD) (*6-9*). Together, these mechanisms are responsible for the rapid and profound lymphopenia observed in patients treated with ATG. However, during or shortly after completion of induction therapy he number of circulating T cells in the blood gradually increases and reverts to normal values (*10*). Interestingly, the recurring T lymphocytes display impaired proliferative capacity even after termination of ATG therapy (*11*). Therefore, additional immunosuppressive mechanisms besides the direct T cell depleting effects are apparent, including masking of antigens, increased clearance of T lymphocytes via the reticuloendothelial system and the expansion of regulatory T cells (T_regs_) (*11-14*). Moreover, ATG-mediated effects on the immune system are not restricted to direct actions on T lymphocytes, as previous studies demonstrated an induction of B lymphocyte apoptosis (*15*), immunomodulation of natural killer cells (*16*) and inhibition of *in vitro* monocytes-derived dendritic cell maturation (*17, 18*). However, the underlying mechanisms responsible for the latter effect are not fully understood.

Immunological responses in T cells requires the engagement of the T cell receptor (TCR) together with an presented antigen and a second co-stimulatory signal (*19*). Depending on the co-stimulatory signal T-cell effector functions are either promoted or dampened (*20*). Continuous stimulation of the TCR, due to ongoing antigen presentation, as seen after solid organ transplantation, has been shown to upregulate the inhibitory co-receptor programmed cell death protein 1 (PD-1) on the surface of activated T cells (*21*). Ligation of this receptor by its ligand programmed death receptor ligand 1 (PDL-1) transduces an inhibitory signal in activated T lymphocytes and decreases their effector functions including proliferation and cytokine release (*22*). Several studies have already reported on the importance of the PD1:PDL-1 axis for maintenance of autoimmunity and peripheral tolerance after solid organ transplantation (*23-25*). However, an involvement of the PD1:PDL-1 system in ATG-induced immunosuppression has not been described so far.

In this study we examined the transcriptional changes of ATG-treated circulating immune cells from human blood on a single cell level. We demonstrated a novel immunosuppressive action of ATG, by the induction of PDL-1 expression on monocytes. Mechanistically, we were able to show that ATG-mediated release of Interferon-γ (IFN-γ) from T-cells was responsible for the induction of PDL-1 expression on monocytes. The interaction of PDL-1 expressing monocytes with PD1^+^ T-cells led to a reduction of T-cell proliferation and granzyme B production. Together, we identified a novel mode of action by which ATG exerts its immunosuppressive effects, even beyond the already known direct effects on activated T lymphocytes.

## RESULTS

### ATG alters the transcriptional profile of lymphoid and myeloid immune cells from human whole blood

To elucidate effects of ATG beyond T lymphocyte depletion as well as its influences on myeloid immune cells we investigated the transcriptional changes resulting from treatment with ATG. Single cell RNA-sequencing (scRNAseq) of untreated, isotype- and ATG-treated (ATG) human whole blood revealed 11 distinct cell clusters in each of the investigated conditions (Fig. 1A). Cell types were assigned to clusters based on the expression of established marker genes (Fig S1A). We identified different subsets of *CD4*^*+*^ T cells (naive *CD4*^*+*^ T cells, effector *CD4*^*+*^ T cells, Tregs), *CD8*^*+*^ T cells (naive *CD8*^*+*^ T cells, effector *CD8*^*+*^ T cells 1, effector *CD8*^*+*^ T cells 2) as well as NK/T cells, B cells, monocytes, *FCGR3A*^*+*^ monocytes and dendritic cells. The relative cluster frequencies across all conditions were highly similar, with the exception of naive *CD4*^*+*^ T cells which accounted for 41.81% of all captured cells in ATG as opposed to 26.14% and 28.18% in untreated and isotype, respectively (Fig. 1A). ATG altered gene expression in all cell types relative to isotype-treated control (Fig. S1B). In the lymphoid subsets we were particularly interested in influences of ATG on cell types with cytolytic effector functions. Effector *CD8*^*+*^ T cells 1 (104 upregulated, 64 downregulated), effector *CD8*^*+*^ T cells 2 (39 upregulated, 43 downregulated) and NK/T cells (116 upregulated, 105 downregulated) showed considerable changes after treatment with ATG. As they hold high functional resemblance, we next determined the overlapping transcriptional regulations that were induced by ATG in these cell types. Between them 31 genes were commonly upregulated by ATG (Fig. S1C), while 29 were downregulated (Fig. S1D). Amongst the upregulated genes we identified several members of the chemokine gene family (*CCL2, CCL3, CCL3L1, CCL4, CCL8, CXCL8, XCL1* and *XCL2*), the TNF-receptor superfamily (*TNFRSF4* and *TNFRSF9*) and *IFNG* (Fig. S1C). Interestingly, the myeloid cell types captured in our analysis were also strongly affected by treatment with ATG resulting in an upregulation of 173, 119 and 76 genes, whereas 97, 77 and 40 were downregulated (Fig. S1B) in ATG-treated monocytes, *FCGR3A*^*+*^ monocytes and Dendritic cells, respectively. Amongst the 33 genes that were commonly upregulated by ATG in these myeloid cell types we identified the IFN-γ response chemokines *CXCL9, CXCL10, CXCL11* as well as *CD274* (Fig. S1E), which encodes for programmed death-ligand 1 (PDL-1) (*26*). Monocytes were the most abundant myeloid cell population captured in our analysis and revealed the most transcriptional changes (Fig. 1A and Fig. S1B). Significantly regulated genes with the highest log_2_ fold changes in monocytes treated with ATG compared to isotype are shown in Fig. 1B, while volcano plots highlighting the regulations in the other cell types are provided as supplementary information (Fig. S2). An overrepresentation analysis for immune system processes affected by genes upregulated in monocytes of ATG treated whole blood revealed significant associations with cellular responses to interferon-γ, immune cell chemotaxis, antibody-dependent cellular cytotoxicity and the regulation of CD8^+^ T cell activation (Fig. 1C). Genes downregulated in ATG-treated monocytes were associated with immune system processes involved in macrophage activation and migration amongst others (Fig. 1D). In line with the transcriptional alterations, analysis of the plasma of ATG treated whole blood showed increased protein release of several cytokines and chemokines. In total we detected 12 factors that were significantly increased in the plasma of ATG-treated whole blood compared to isotype-treated controls (Fig. 1E), including members of the CCL- or CXCL-family, IL-2, TNF-α, GM-CSF and IFN-γ. Similar findings were observed in conditioned media (CM) of purified PBMCs treated with ATG where the increase of CCLs, CXCLs and other cytokines, including IFN-γ, was even more pronounced (Fig. S3A).

**Fig. 1.**
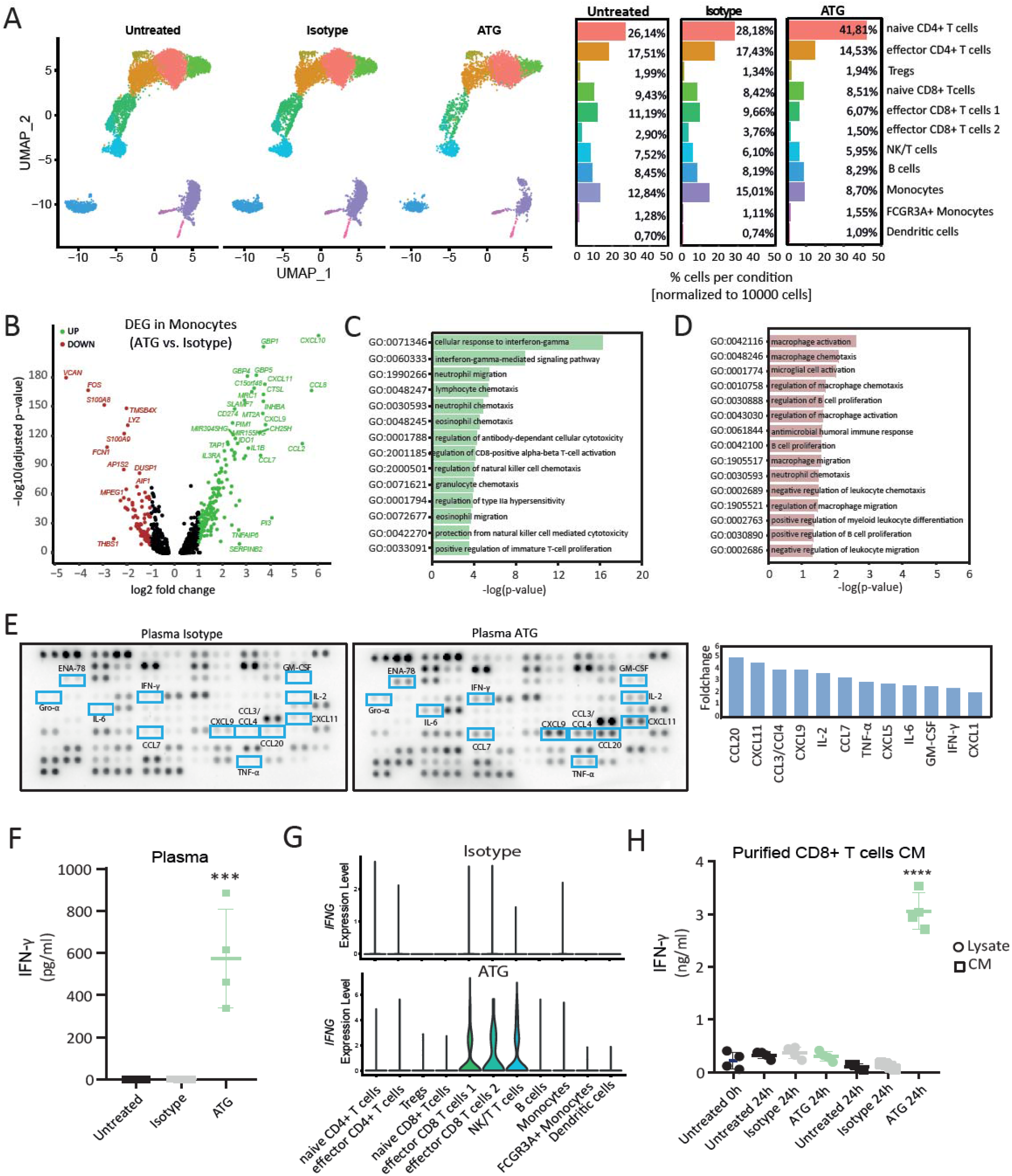
Single cell RNA-sequencing of ATG-treated white blood cells reveals transcriptional alterations in monocytes in response to IFN-γ. **(A)** Uniform Manifold Approximation and Projection (UMAP) plot of untreated (Untreated), isotype-treated (Isotype) and ATG-treated (ATG) immune cells from whole blood reveals 11 distinct cell types present in all conditions. Relative distribution of each cell type across all investigated conditions is shown in the barplots. N= 2 donors per condition (**B**) Volcano plot showing differentially up-(green, log_2_FC >1 and adj. p-value < 0.05) and downregulated (red, log_2_FC < -1 and adj. p-value < 0.05) genes (DEG) in monocytes (ATG vs. Isotype) with annotations. Adj. p-values were obtained after Benjamini-Hochberg correction. Bar plot showing significantly overrepresented Immune System Processes associated with **(C)** up- and **(D)** downregulated genes in monocytes (ATG vs. Isotype). **(E)** Immunodetection array membrane of plasma of whole blood treated with ATG and isotype control. Pooled supernatants of 3 donors per condition were analyzed. Proteins with a > 2-fold increase are shown in blue brackets and highlighted in the bar graph. (**F**) Quantification of IFN-γ by ELISA in plasma from untreated (black), isotype-treated (grey) and ATG-treated (green) whole blood. N = 4 donors. Asterisk denotes p-value = 0.0005. Ordinary one-way ANOVA was performed followed by Dennett’s multiple comparisons. **(G)** Expression of *IFNG* across all clusters in Isotype and ATG. Grade of gene expression is indicated by violin plot height while width represents proportion of positive cells. **(H)** Quantification of IFN-γ by ELISA in lysates (circle) and conditioned medium (square) of untreated (black), isotype-treated (grey) and ATG-treated (green) purified CD8^+^ T cells. N = 4 donors. Asterisk denotes p-value < 0.0001. Ordinary one-way ANOVA was performed followed by Dennett’s multiple comparisons.

### ATG induces IFN-γ production and release in CD8^+^ effector T cell subsets

As “cellular response to interferon-gamma” was the strongest regulated GO-term and several of the identified upregulated cyto- and chemokines were IFN-*γ* inducible, we next investigated the regulation and action of IFN-*γ* after ATG stimulation in more detail. First, we corroborated our finding from the proteome profiler with an IFN-*γ* ELISA and found that absolute plasma levels were significantly increased after treatment with ATG (575.5 ± 235.1 pg/ml; p-value = 0.0005) compared to isotype and untreated control (both below the assays detection limit, Fig. 1F). Protein levels of IFN-γ in conditioned media of isolated PBMCs were also significantly higher in ATG-treated cells than in controls (Untreated: 22.76 ± 22.76 pg/ml; Isotype: below assay detection limit; ATG: 575.0 ± 101.3 pg/ml, p-value = 0.0008) (Fig. S3B). Only the subsets of effector *CD8*^*+*^ T cells as well as the NK/T cells showed increased expression of Interferon-Gamma mRNA (*IFNG*) following treatment with ATG (Fig 1G). Next we aimed to further corroborate the IFN-γ modulating effects of ATG on CD8^+^ T cells. Conditioned media of ATG-treated CD8^+^ T cells revealed significantly higher levels for IFN-γ (3059 ± 173 pg/ml; p-value < 0.0001) as compared to isotype (122 ± 47 pg/ml) and untreated controls (98 ± 28 pg/ml) after purification and stimulation (Fig. 1H). Moreover, measurement of IFN-*γ* in CD8^+^ T cell lysate immediately after isolation (218 ± 80 pg/ml) as well as 24 hours after stimulation (untreated 24h: 326 ± 35 pg/ml; isotype 24h: 367 ± 56 pg/ml; ATG 24h: 301 ± 45 pg/ml) showed no upregulation of IFN-γ levels by ATG, suggesting that newly produced IFN-*γ* is immediately released by the cells (Fig.1H). Together our data indicate that ATG induces the production of IFN-γ in cytotoxic CD8^+^ T cells which in turn affects monocyte function.

### Surface expression of PDL-1 is increased on monocytes of ATG-treated whole blood

PDL-1, encoded by CD274, is known for its involvement in the regulation of T-cell activity via engagement of PD-1 (*27*). In addition to several chemokines and cytokines, *CD274* was also significantly upregulated by ATG in monocytes, *FCGR3A*^*+*^ monocytes and dendritic cells from whole blood (Fig. 2A). Treatment with ATG resulted in a more than 2.5-log_2_fold upregulation of CD274 expression in monocytes (Fig. 2A). Almost 60% of all monocytes showed high levels of CD274 after ATG stimulation (0.32 ± 0.32 % in untreated, 1.96 ± 1.613% in isotype treated and 58.74 ± 10.01% in ATG treated monocytes) (Fig. 2B). These changes were further affirmed by assessment of surface protein expression of PDL-1 in monocytes treated with ATG (Fig. 3C). 24 hours after stimulation of whole blood with ATG 89.68 ± 5.2% (p-value < 0.0001) of monocytes were positive for PDL-1 while untreated (4.6 ± 0.5%) and isotype (6.6 ± 0.52%) treated samples showed little if any PDL-1 expression (Fig. 2C). Similarly, 88.9 ± 2.35% (p-value: <0.0001) of monocytes in ATG-treated purified PBMCs were positive for PDL-1 as opposed to 2.105 ± 0.46% in untreated and 4.932 ± 0.76% in isotype controls, respectively (Fig. 2C). Together these data suggest, that ATG potently upregulates the expression of *CD274* on monocytes which translates to an increase of surface expression of PDL-1 on monocytes from whole blood and purified PBMCs.

**Fig. 2.**
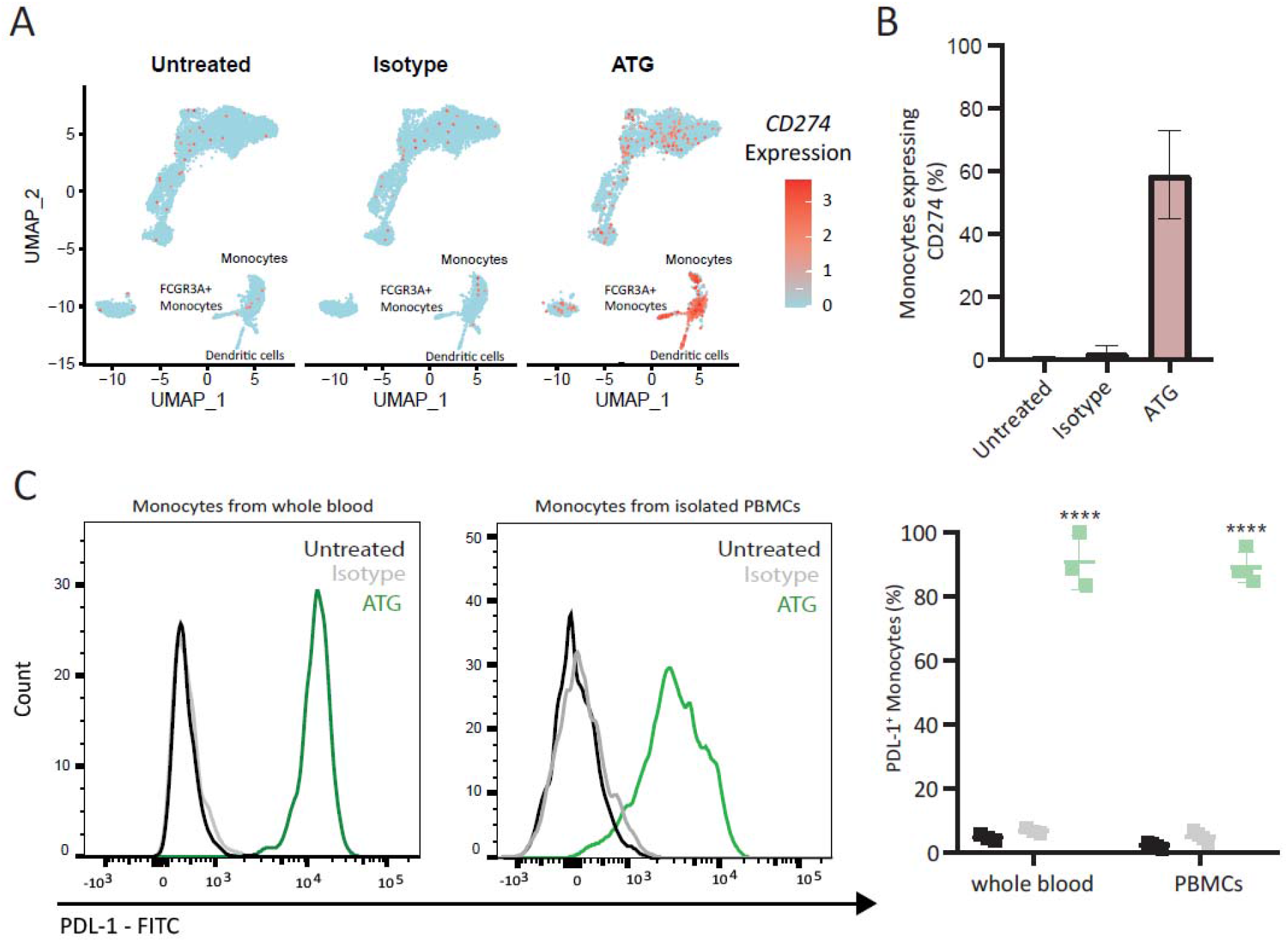
ATG upregulated *CD274* in monocytes from whole blood *ex vivo*. (**A**) Feature plot for *CD274* of untreated, isotype- and ATG-treated white blood cells. Monocytes, *FCGR3A*+ Monocytes and Dendritic cells are annotated and show higher expression of *CD274* in ATG than in controls. (**B**) Bar plot showing the mean percentage of Monocytes expressing *CD274* and standard error of the mean (SEM) in Untreated, Isotype and ATG. N = 2 donors. (**C**) Flow cytometry analysis for PDL-1 of ATG treated whole blood and purified PBMCs. Histogram shows Monocytes of Untreated (black), Isotype (grey) and ATG (green) from whole blood and purified PBMCs. Significantly higher percentages of PDL-1+- monocytes are detected in ATG when compared to Untreated and Isotype, respectively. ** indicates p-value < 0.01; **** indicates p-value < 0.0001

**Fig. 3.**
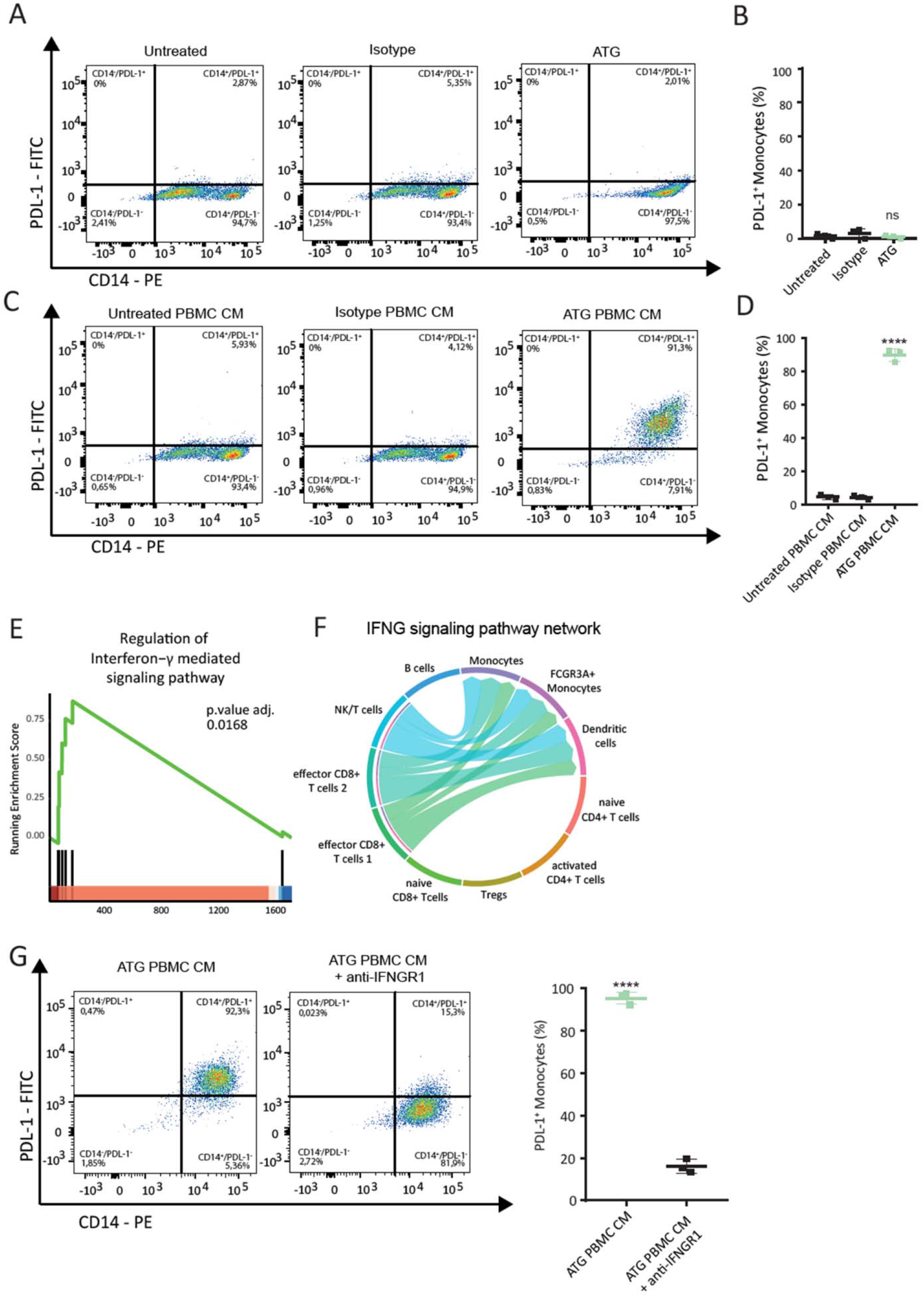
IFN-γ modulates upregulation of PDL-1 on purified monocytes. (**A**) Representative FACS-plots of purified monocytes assessed for surface expression of PDL-1 24 hour after treatment with isotype antibody, ATG, conditioned medium of untreated, isotype-treated and ATG-treated PBMCs. Untreated monocytes served as control. PDL1^+^-monocytes are detected in the upper right quadrant. (**B**) No statistically significant differences were observed in purified monocytes treated directly with ATG. (**C & D**) Conditioned medium of ATG-treated PBMCs significantly increase PDL-1 expression on purified monocytes displaying a p-value < 0.0001 (**E**) GSEA enrichment plot for representative signaling pathway enriched in monocytes from ATG-treated whole blood. Green line depicts running enrichment score, black vertical lines indicate rank of genes responsible for enrichment and position in list of DEG. (**F**) Chord diagram of *IFNG* signaling pathway network connecting effector *CD8+* T cell subsets and NK/T cells with monocytes, *FCGR3A+* monocytes and dendritic cells. (**G**) Representative FACS plots of purified monocytes treated with conditioned media of ATG treated PBMCs. Surface expression of PDL-1 was assessed with prior inhibition of IFN-γR1 and without. Asterisk denotes p-value < 0.0001 and was calculated by one-tailed student’s t-test.

### IFN-γ modulates surface expression of PDL-1 on purified monocytes

To identify the mechanism by which ATG increases PDL-1 expression in monocytes, we first investigated whether ATG-induced PDL-1 expression is a result of a direct or indirect stimulation of monocytes. For that reason, we purified monocytes from whole blood and stimulated them with ATG. Purity of isolated monocytes was assessed by flow cytometry and was above 95% (Fig. S4A). Interestingly, direct stimulation of purified CD14^+^-monocytes with ATG did not lead to an upregulation of PDL-1-expression (Fig. 3A and B). Since our transcriptional data identified strong regulation of several cytokines, we next investigated whether the modulation of PDL-1 was dependent on soluble factors induced by ATG. We therefore stimulated PBMCs with ATG for 24 hours and added the resulting supernatant on purified monocytes. Indeed, monocytes incubated with conditioned media of ATG-treated PBMCs were strongly positive for PDL-1, while monocytes treated with conditioned media of untreated PBMCs showed almost no PDL-1 (Fig. 3C and D; untreated PBMC SN: 4.44 ± 0.85%; isotype PBMC CM: 4.12 ± 0.78%; ATG PBMC CM: 89.9 ± 2.07%; p-value < 0.0001, for gating strategy see Fig. S4B). Furthermore, we evaluated the long-term prevalence of PDL-1^+^ - monocytes induced by the conditioned medium of ATG-treated PBMCs. Significantly higher mean percentages of PDL-1^+^ monocytes were detected following treatment with ATG CM compared to controls for up to 120 hours (Fig. S5). To investigate the signaling pathway(s) responsible for the observed effect, we next performed gene set enrichment analysis of genes regulated in Monocytes by ATG. This analysis showed a significant enrichment of genes regulated by interferon-γ (Fig. 3E and. S6) and corroborated the top 2 immune system processes detected in our overrepresentation analysis (Fig. 1C). In addition, we observed a significant regulation of the IFN-γ signaling pathway network between effector *CD8*^*+*^ T-cell subsets and myeloid cell types, as shown by the chord diagram (Fig. 3F), suggesting a ligand-receptor communication based on the interactions of *IFNG* with its receptors *IFNGR1* and *IFNGR2*. To validate our bioinformatics data, we investigated the expression and function of the IFN-γ - receptors on monocytes. Flow cytometry analysis confirmed the expression of IFN-γR1 and IFN-γR2 on purified monocytes (Fig. S4C). Addition of IFN-γR1 but not IFN-γR2 blocking antibodies almost completely abolished PDL-1^+^-upregulation in monocytes after treatment with PBMC-conditioned media (95.4 ± 1.58% vs. 16.07 ± 1.86%; p-value < 0.0001) (Fig. 3H and S7). These findings suggest that engagement of IFNγR1 by IFN-γ is required to induce PDL-1 on purified monocytes.

### Inhibition of STAT1 by ruxolitinib abolishes induction of PDL-1 on monocytes by paracrine factors

Since IFN-γR1-signalling is known to involve the activation of the JAK/STAT pathway (*28*), we next investigated whether activation of STAT1 influences PDL-1 expression on monocytes. *STAT1* mRNA expression was upregulated in monocytes, *FCGR3A*^*+*^ monocytes and dendritic cells (Fig. S1E). Moreover, we detected a significant enrichment of the receptor signaling pathway via JAK/STAT (adjusted p-value: 0.0022) in monocytes of ATG-treated whole blood (Fig. 4A). Further investigation of the JAK/STAT gene family members, involved in the key enrichment of this pathway, revealed strong upregulation of *JAK2, STAT1, STAT2 and STAT3* in monocytes (Fig. 4B), but also other cell types of ATG-treated whole blood (Fig. S8). In contrast to the controls, treatment with conditioned media of ATG-treated PBMCs increased phosphorylation of STAT1 in purified monocytes (Fig. 4C). Pre-incubation of purified monocytes with the JAK-inhibitor ruxolitinib strongly reduced this effect (Fig. 4C). Importantly, addition of ruxolitinib to monocytes treated with conditioned media of ATG-treated PBMCs strongly inhibited PDL-1^+^-expression (Fig. 4D). While 92.23 ± 3.06% of monocytes incubated with ATG PBMC CM were positive for PDL-1 only 4.97 ± 1.84% of monocytes were positive for PDL-1 when cells were pre-treated with ruxolitinib (Fig. 4E, p-value < 0.0001). Taken together, we show that modulation of PDL-1 surface expression on monocytes by ATG is regulated by IFN-γ and dependent on its binding to IFN-γR1 and downstream activation of the JAK/STAT signaling pathway.

**Fig. 4.**
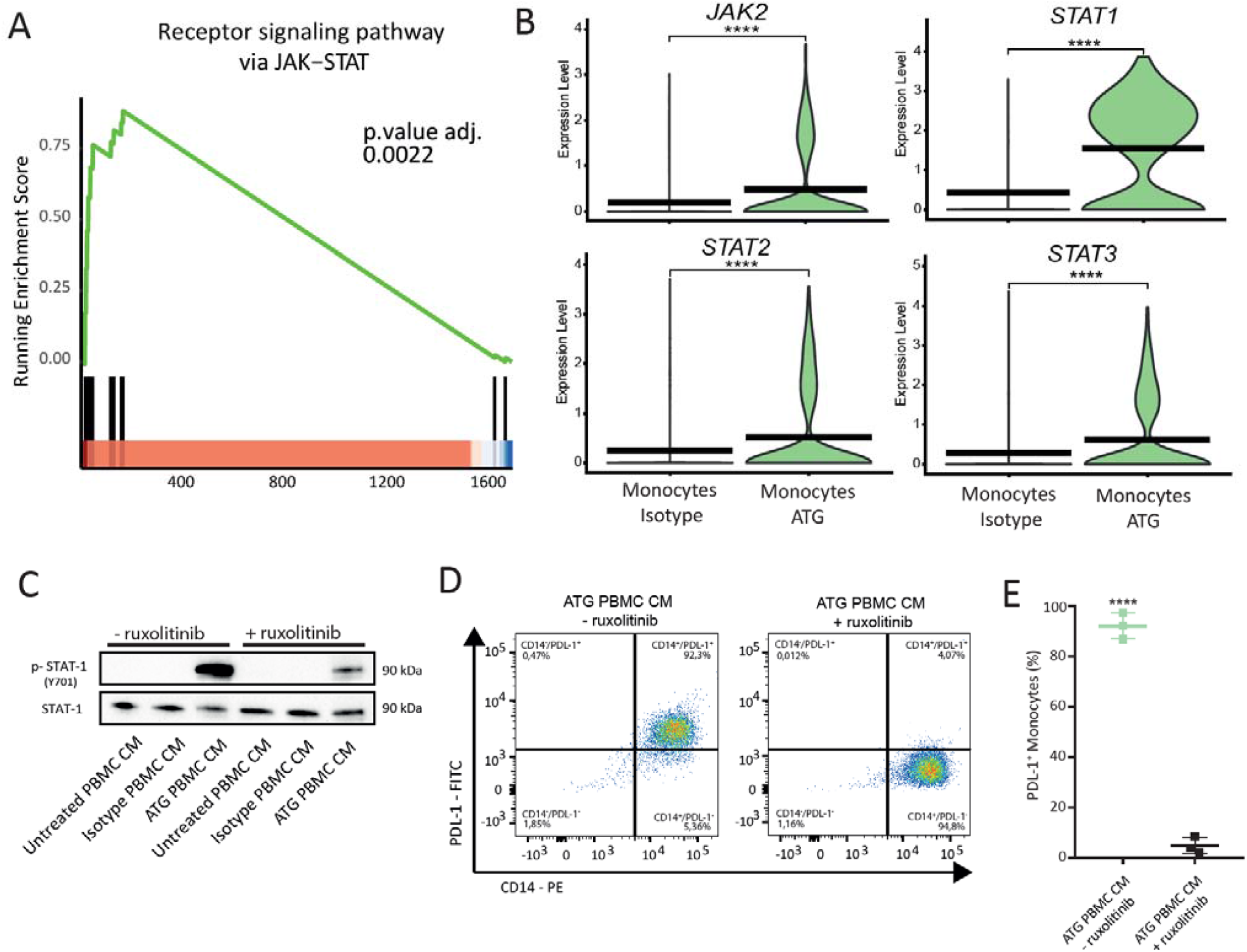
STAT-1 activation is required for induction of PDL-1 on purified monocytes. (**A**) GSEA enrichment plot for receptor signaling pathway via JAK-STAT in monocytes from ATG-treated whole blood. Green line depicts running enrichment score, black vertical lines indicate genes responsible for enrichment and position in ranked list of DEG. (**B**) Violin plots depicting differential expression *JAK2, STAT1, STAT2* and *STAT3* in monocytes following stimulation with ATG. Expression levels are indicated by violin plot height while width represents proportion of positive cells. Crossbars mark mean expression. **** indicate p-value < 0.0001 (**C**) Representative western blots of purified monocytes pretreated with the JAK/STAT inhibitor ruxolitinib for 1 hour before stimulation with conditioned media of ATG treated PBMCs and controls. Total STAT1 and phosphorylated STAT1 were assessed after 2 hours. (**E**) FACS plots showing the percentage of monocytes positive for PDL-1 24 hours after addition of conditioned medium of ATG-treated PBMCs without and with prior addition of ruxolitinib. N=3 donors. Asterisk denotes p-value < 0.0001. Student’s t-test was used to determine significance.

### PDL1^+^-monocytes inhibit CD8^+^-T cell proliferation and release of granzyme B *in vitro*

Activation of co-stimulatory and co-inhibitory signals influences effector functions of T lymphocytes (*29*). The binding of PDL-1 to PD-1 confers an inhibitory signal (*30*). Monocytes in the ATG condition displayed a positive enrichment for biological processes associated with the regulation of CD8^+^ T cell activation (adjusted p-value: 0.0124), negative regulation of T cell proliferation (adjusted p-value: 0.0086) and tolerance induction (adjusted p-value: 0.0086) (Fig. 5A). *CD274, IDO1, IRF1, HLA-A* and *HLA-E* were identified as core enrichment genes for these processes (Fig. 5B). Therefore, we thought to assess whether PDL-1^+^ monocytes are able to suppress effector functions of activated CD8^*+*^ T cells *in vitro*. Co-culture of PDL-1^+^-monocytes with αCD3/αCD28 activated CD8^+^ T cells significantly reduced their proliferative capacity (Fig. 5C, for gating strategy see Fig. S4D). When incubated with untreated control monocytes 55.03% ± 2.37% of CD8^+^ T had divided ≤ 3 and 44.98% ± 2.37% ≥ 4 times after 5 days (Fig. 5C and D). In contrast, 81.63% ± 2.98% of activated CD8^+^ T cells had divided ≤ 3 times and18.38 ± 2.98% ≥ 4 times (Fig 5D; p-value = 0.0009) after co-culture with PDL-1^+^ monocytes. This effect was significantly less pronounced when PDL1^+^ monocytes were pre-treated with durvalumab, a monoclonal antibody directed against PDL-1, prior to co-culture with activated T lymphocytes (59.78 ± 4.44% displayed ≤ 3 and 40.23 ± 4.44% ≥ 4 divisions (Fig. 5D; p-value = 0.0034).

**Fig. 5.**
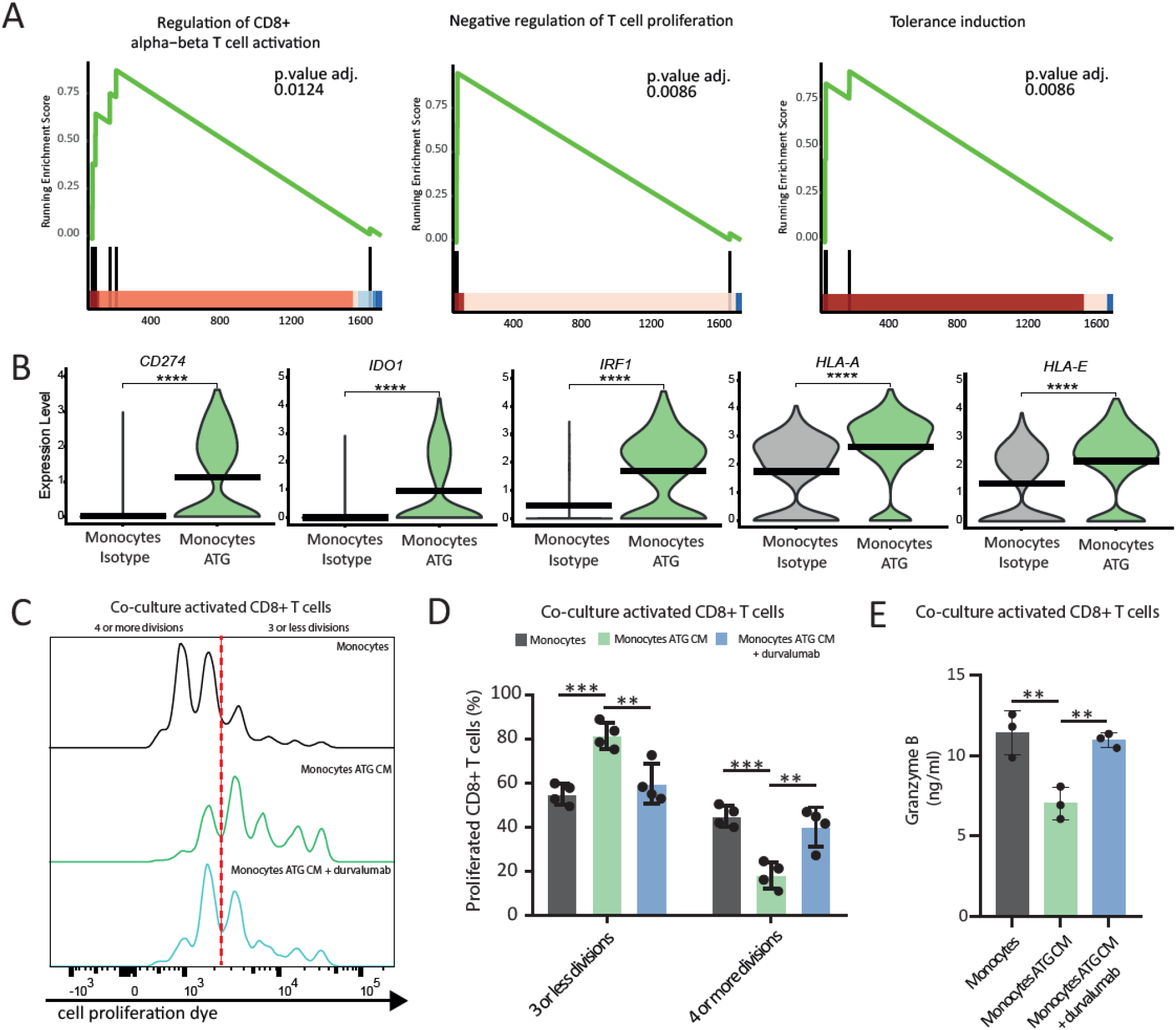
PDL-1^+^ monocytes reduce proliferation and granzyme B release of activated CD8^+^ T cells. **(A)** GSEA enrichment plot for regulations of T-cell functions and tolerance induction in monocytes following treatment with ATG. Green lines depict running enrichment score, black vertical lines indicate genes responsible for enrichment and position in ranked list of DEG. **(B)** Violin plots of core enrichment genes *CD274, IDO1, IRF1, HLA-A* and *HLA-E* in monocytes isotype versus monocytes ATG. Expression levels are indicated by violin plot height while width represents proportion of positive cells. Crossbars mark mean expression. **** indicate p-value < 0.0001. **(C)** Representative histogram of proliferative capacity of activated CD8^+^ T cells co-cultured with monocytes (black), PDL-1^+^ monocytes (green) and durvalumab pre-treated PDL1^+^ monocytes (blue). Non-proliferating (furthest right) and proliferating cell populations are reflected by intensity of cell proliferation dye staining. Dashed red line separates cells with 3 or less divisions from cells with 4 or more divisions. **(D)** Bar graph depicting mean percentages of CD8^+^ T cells with 3 or less proliferations and 4 or more proliferations at day 5. N = 4 donors per group. *** p-value = 0.0009; ** p-value = 0.0034. One-way ANOVA with Dennett’s multiple comparison was used to determine statistically significant differences. **(E)** Concentration of granzyme B in conditioned medium of CD8^+^ T cells co-cultured with monocytes assessed by ELISA. ** p-value < 0.01.

Next, we also measured protein levels of the effector cytokine granzyme B in supernatants of activated CD8^+^ T cells co-cultured with monocytes. Addition of PDL-1^+^ monocytes resulted in significantly lower levels of granzyme B (7.01 ± 0.58 ng/ml) when compared to untreated controls (11.41 ng/ml ± 0.79; p-value = 0.0046) and anti-PDL-1 antibody pre-treated PDL-1^+^-monocytes (10.96 ± 0.26 ng/ml; p-value = 0.0077) (Fig. 5E). Together, these data show that PDL-1^+^-monocytes inhibit effector functions of activated CD8^+^-T cells, resulting in reduced T-cell proliferation and granzyme B secretion.

## DISCUSSION

The immunosuppressive action of ATG is generally attributed to its T cell depleting properties (*7, 9, 13*). However, increasing evidence from several groups suggests additional modes of action, mediated either directly on distinct T lymphocyte subsets, or indirectly via the release of paracrine factors (*12, 31-33*). Using scRNAseq we detected numerous alterations in gene expression across lymphoid and myeloid cell types of white blood cells from human whole blood treated with ATG. Notably, classical monocytes displayed significant changes that were tightly connected to responses to IFN-γ. As a result, surface expression of PDL-1 was increased in these cells, enabling them to functionally impair the proliferative capacity and release of granzyme B of activated CD8^+^ T cells, representing an additional mechanism by which ATG exerts its immunosuppressive actions.

Here we provide a broad overview on the transcriptional changes modulated by *ex vivo* treatment of human whole blood with ATG capturing different subsets of lymphoid and myeloid cell types. Transcriptional changes in the lymphoid cells are explainable by the antigen specificity of ATG (*34*). Influences on myeloid cell types such as natural killer cells and monocyte-derived dendritic cells have been previously reported by others (*17, 32*). Dalle *et al*. showed increased production of IFN-γ in natural killer cells (*32*), while Roider *et al*. reported on the *in vitro* induction of tolerogenic dendritic cells by ATG (*31*). We and others could previously show that PBMCs treated with ATG show increased production of IFN-γ (*35, 36*). In line with these reports, our analysis revealed an upregulation of *IFNG* in NK/T cells after stimulation with ATG. Beyond that, we observed increased expression of *IFNG* in the subsets of effector *CD8*^*+*^ T cells following treatment with ATG and identified these to be specifically modulated by ATG to increase production and release of IFN-γ. While IFN-γ is generally considered as a pro-inflammatory cytokine several reports suggest immunosuppressive roles for IFN-γ (*37*). Priming of human mesenchymal stem cells with IFN-γ modulated their immunosuppressive capacity and resulted in clinical improvement and prolonged survival in a murine model of Graft versus host disease (*38*). Furthermore, IFN-γ has been shown to confer immunosuppressive actions by modulating expression of the inhibitory immune-receptor ligand PDL-1 on various cell types (*39-41*). In addition to IFN-γ numerous other cyto-and chemokines were induced by ATG including particularly members of the CCL- and CXCL-family as well as immunomodulatory cytokines. Especially CXCL9, CXCL10 and CXCL11 have been considered as IFN-γ response cytokines and reported to modulate the expression of PDL-1 in gastric cancer cells (*42*). Their presence might explain why after inhibition of IFN-γR1 the induction of PDL-1 on monocytes was not completely abolished. Furthermore, these factors bind to the chemokine receptor CXCR3 and transduce their signal via activation of the JAK/STAT pathways (*43*), supporting the superior inhibition of PDL-1 induction by the JAK/STAT inhibitor used in our experiments. The overrepresentation of paracrine factors regulated by ATG and the resulting activation of intracellular signaling pathways highlight their relevance for the effects mediated by ATG. Our data further substantiate the importance of paracrine factors released by white blood cells in response to ATG and describe an additional immunosuppressive mechanism beyond the already known depletion of T-cells.

The observed increase in surface expression of PDL-1 on monocytes was mediated by paracrine signaling of IFN-γ and lead to an inhibitory effect on activated CD8^+^-T cells. This mechanism has previously been described as an evasion mechanism of different tumors entities (*44, 45*). Binding of IFN-γ to IFN-γR1 modulated expression of PDL-1 on tumor cells and suppressed anti-tumor host immunity leading to disease progression, increased dissemination and worse overall outcome (*46*).While these regulations are clearly detrimental in oncologic settings, following transplantation modulation of the PD1:PDL-1 axis can significantly improve graft survival and peripheral tolerance (*47*). Pre-clinical work by Borges *et al*. revealed a significant increase in graft tolerance of PD-1 overexpressing T cells in a fully MHC-mismatched murine cardiac transplantation model (*48*). Interestingly, transplantation of PDL-1-knockout donor hearts into PD-1 overexpressing mice resulted in immediate graft rejections strongly underling the importance of the receptor-ligand-pair for peripheral tolerance (*48*). Moreover, expression of PDL-1 can be found on other cell types besides myeloid cells. Recently, Bracamonte-Baran *et al*. reported on the increased presence of PDL-1 on graft endothelial cells and their inverse correlation with infiltration of CD8^+^ T lymphocytes in myocardial biopsies of heart transplant recipients (*49*). Furthermore, they show that in fully MHC-mismatched mice a conditional knockout of PDL-1 in graft endothelial cells significantly decreased graft survival when compared to PDL-1 expressing controls (*49*). Our observations are restricted to immune cells from whole blood. Still, ATG and paracrine factors induced by ATG might also affect other cell types found in the vasculature or even in the transplanted organ and similarly modulate PDL-1 or alternative pathways to contribute to graft tolerance by reducing T-cell mediated tissue damages. The emergence of immune-therapeutic drugs for the targeted inhibition of co-stimulatory signals with inhibitory effects on activation and proliferation of T-cells, i.e. PDL-1 and PD-1 amongst others, has advanced the treatment of patients affected by oncologic malignancies (*50*). However, severe adverse effects related to excessive activation of the immune system by therapy with immune-checkpoint inhibition have been reported in these patients (*51*). These immune-related adverse events most commonly involve hepatitis, colitis, myocarditis and dysregulations of endocrine systems including the pancreas and adrenal glands (*52*). Interestingly, several reports showed that ATG is able to counteract adverse events caused by immune checkpoint inhibition (*53-55*). While this effect is in part explainable by its T-cell depleting properties, based on our data, it is tempting to speculate that induction of PDL-1 on monocytes, or other cell types, potentially contributes to the inhibition of autoreactive T-cells, resulting in an amelioration of the reported adverse effects.

Moreover, transplanted patients naturally are at a higher risk to develop neoplasia due to ongoing maintenance immunosuppressive therapy (*56, 57*). While there are no large randomized clinical trials available, several case reports described a high risk of graft failure and rejection in solid organ transplant recipients treated with immune checkpoint inhibitors underlining the importance of the PD1:PDL-1 axis for maintenance of peripheral graft tolerance (*58, 59*). Whether ATG might prove beneficial in these specific settings where fine margins between maintenance of graft survival are confronted with the requirement of anti-tumor immunity has to be addressed in future studies. Our data show that ATG is able to increase levels of PDL-1 on monocytes and other myeloid blood cells and the interaction of these cells with activated T cells results in the reduction of their effector functions including proliferation and release of granzyme B.

Our findings are limited to pre-clinical *ex vivo* and *in vitro* data. Studies addressing the role of ATG induced PDL-1 on myeloid cells and its relevance in patient settings will be needed to clarify to which degree this effect contributes to ATG-mediated graft tolerance. In addition, such studies will be better suited to investigate influences of repeated administrations of ATG, as performed in standardized induction protocols, and enable a prolonged observation of the described effects. Nevertheless, our long-term studies already showed a prolonged positivity of PDL-1 on monocytes treated with a single administration of conditioned medium of ATG-treated PBMCs. In addition, immunosuppressive induction therapy in patients with high immunological risk of graft rejections usually includes additional drugs such as cyclosporine A, mycophenolate-mofetil and systemic glucocorticoids (*60*). Of note, pretreatment of PBMCs with an immunosuppressive dose of hydrocortisone prior to addition of ATG did not decrease PDL-1 on monocytes (Fig. S9). This suggests that combined use of these agents does not interfere with PDL-1 induction. Further studies are needed to reliably dissect potential interactions between these and other compounds regularly used for immunosuppression in combination with ATG.

In conclusion, our study provides a comprehensive overview on the transcriptional changes in immune cells from human whole blood treated with ATG on a single cellular level. We identify an increase in surface expression of PDL-1 on monocytes as a result of paracrine signaling via IFN-γ/STAT/JAK. Additionally, we show that PDL-1^+^ monocytes potently inhibit effector functions of activated CD8^+^ T cells *in vitro*. Altogether, our data suggest an additional mechanism by which ATG indirectly modulates immunosuppressive actions on cell-mediated rejections in white blood cells.

## MATERIALS AND METHODS

### Study Design

The aim of this study was to investigate the changes induced by *ex vivo* treatment of human whole blood with ATG and uncover potential novel mechanisms by which ATG regulates T-lymphocyte functions. To this end, we used single cell RNA-sequencing to identify transcriptional alterations set in motion by ATG in different immunological subsets of lymphoid and myeloid white blood cells. For scRNA-seq 2 donors were sampled per experimental group.

To verify our bioinformatics data, we performed a series of cell culture and flow cytometry based assays. For the *in vitro* experiments involving purified PBMCs and purified monocytes the number of individual replicates as well as statistically significant differences were shown in the figure legends and included at least three donors per experiment. We examined the effects of ATG-mediated paracrine factors on purified monocytes and identified a crucial role of IFN-*γ* in the induction of the immune-regulatory co-receptor PDL-1. We also delineated the functional contributions of PDL-1^+^-monocytes on activated CD8+ T-cells *in vitro*.

### Ethics statement

This study was conducted in accordance with the Declaration of Helsinki and local regulations. Blood samples were obtained from healthy volunteers who had given their consent to donate prior to participation. The Institutional Review Board of the Medical University of Vienna approved this study (Ethics committee votes: 1539/2017). All donors provided written informed consent.

### Preparation of single cell suspension of human whole blood

For scRNAseq, heparinized human whole blood was drawn from two age-matched male donors. A total of 3 ml of whole blood was either treated with 100µg/ml ATG (Grafalon, Neovii Biotech GmbH, Gräfelfing, Germany), an equivalent dose of polyclonal rabbit isotype control (ab37415, Abcam, Cambridge, UK) or left untreated. The used dosage corresponds to the blood levels detected in patients undergoing ATG induction (*61*). Samples were incubated at 37°C for 8h. Next, red blood cells were removed by Red Blood Cell Lysis Buffer (Abcam). Cells were then washed twice with PBS containing 0.04% bovine serum albumin (BSA, Gibco) and sequentially passed through 100- and 40-µm cell strainer. The LUNA-FL Dual Fluorescence Cell Counter (BioCat, Heidelberg, Germany) and the Acridine Orange/Propidium Iodide Cell Viability Kit (Logos Biosystems, Gyeonggi-do, South Korea) were used to adjust cell count to 1×10^6^ cells/ml with a viability above 90%.

### Gel Bead-in Emulsion (GEMs) - and library preparation

Single-cell RNA-seq was performed using the 10X Genomics Chromium Single Cell Controller (10X Genomics, Pleasanton, CA, USA) with the Chromium Single Cell 3′ V3 Kit following manufacturer’s instructions. After quality control, sequencing was performed by the Biomedical Sequencing Core Facility of the Center for Molecular Medicine (Center for Molecular Medicine, Vienna, Austria) on an Illumina HiSeq 3000/4000 (Illumina, San Diego, CA, USA). For donor 1, we detected 2094 cells in total, while for donor 2 altogether 18257 cells were captured. Raw sequencing data were then processed with the Cell Ranger v3.0.2 software (10X Genomics, Pleasanton, CA, USA) for demultiplexing and alignment to a reference genome (GRCh38).

### Single cell RNA-sequencing data analysis

Secondary data analysis was performed using R Studio in R (The R Foundation, Vienna, Austria) using the R software package “Seurat” (Seurat v.4.0.0, Satija Lab, New York, NY, USA). Cells were first analyzed for their unique molecular identifiers (UMI) and mitochondrial gene counts to remove unwanted variations in the scRNA-seq data. Cells with Feature RNA counts below 100 or above 2500 and more than 10% of mitochondrial genes were excluded from the data set. Next, we followed the recommended standard workflow for integration of scRNAseq datasets (*62*). Data were scaled and principal component analysis (PCA) was performed. Statistically significant principal components (PCs) were identified by visual inspection using an Elbow plot. Calculation of the Louvain algorithm at a resolution of 0.2 iterations identified a total of 11 communities. The preselected PCs and identified clusters served for Uniform Manifold Approximation and Projection for Dimension Reduction (UMAP). After bioinformatics integration of the datasets of Untreated, Isotype-treated and ATG-treated samples, erythrocytes were removed by excluding all cells with expression of Hemoglobin subunit beta (HBB) > 0.5. Annotation of cell types to the calculated clusters was based on the expression of cell-type-specific marker genes that were determined with the “FindAllMarkers” argument in Seurat. We used UMAP-plots, dot plots, feature plots, volcano plots and violin plots to visualize differences between the investigated conditions. Normalized count numbers were used to determine DEGs. We applied the “FindMarkers” argument using default settings to calculate DEGs for clusters of interest between conditions. A Log2-fold-change increase of gene expression above 1 was considered as upregulation while a decrease below -1 was considered as downregulation. The sets of DEGs were imputed into the Cytoscape (*63*) plug-in ClueGO (*64*) to visualize significantly (p-value < 0.05, kappa score: 0.4) overrepresented Gene Ontologies related to immune system processes for the investigated conditions.

### Gene Set enrichment analysis (GSEA)

GSEA of gene ontologies was calculated in R using the Bioconductor package “clusterprofiler” (*65*). A list of DEGs was calculated comparing every ATG-treated cell type to its matched Isotype control. This list was then sorted in descending order for each cell type. For gene annotation the “org.Hs.eg.db.” package was loaded and executed in R. Non-annotated genes were omitted from further analysis. Using the “gseGO” command we calculated significantly (p-value < 0.05) enriched gene ontologies for biological processes (GO: BP) in the ATG treated cell types. The default settings for minimal and maximal gene set size as well as the number of permutations were maintained for initial analysis. To adjust the calculated p-values and minimize false discovery rate we performed a Benjamini-Hochberg correction. Results were visualized by GSEA plots embedded in the “ggplot2” package.

### CellChat analysis for ligand-receptor interactions

To infer cellular communications between the identified cell types in our scRNA-seq analysis we used the R package “CellChat” (*66*). Firstly, we converted our initial Seurat object into a CellChat object. Next, the CellChat database of receptor and ligand pairs was implemented into the new object. Then we performed an overrepresentation analysis for genes and possible interactions within the CellChat database. The p-value threshold for a significant ligand-receptor interaction was set < 0.05. Cell-cell interactions are displayed as chord diagrams.

### Isolation of PBMCs and cell purification

For in vitro assays PBMCs were isolated using density gradient centrifugation with Ficoll-Paque PLUS (GE Healthcare Bio-Sciences AB, Sweden). Heparinized blood was diluted with phosphate-buffered saline (PBS, Gibco by Life Technologies, Carlsbad, CA, USA) and carefully layered over Ficoll-Paque PLUS. After centrifugation (800g, 15 min, room temperature, with slow acceleration and deceleration), buffy coats containing PBMCs were enriched at the plasma-Ficoll interface. For purification of CD14^+^ monocytes and CD8^+^ T cells we performed magnetic bead separation using CD14 and CD8 magnetic beads (Miltenyi, Bergisch Gladbach, Germany) to enrich cells over the QuadroMACS^™^ Seperator (Miltenyi) according to the manufacturers protocol. Purity of isolated cells was confirmed by flow cytometry using antibodies against CD14-PE (BioLegend, San Diego, California, USA; clone: HCD14) and CD8-PeCy5 (BioLegend; clone: SK1) and ranged from 95 to 99%. A complete list of all antibodies used in this study is provided in supplementary table 1. Acquired cells were counted, diluted to a concentration of 1×10^6^ cells/ml and cultured in RMPI 1640 supplemented with 10% fetal bovine serum (FBS, ThermoFisher Scientific, Waltham, Massachusetts, USA) and 1% Pen/Strep (ThermoFisher Scientific) unless otherwise stated. After 24h of stimulation with ATG PDL-1-FITC (BioLegend, clone: MIH2) antibody was used to assess surface expression on immune cells.

### Proteome Profiler and Enzyme-linked immunosorbent assay

Whole blood samples and isolated PBMCs were treated with ATG (100µg/ml) or Isotype control (100µl/ml) and cultured in standard cell culture conditions for 24 hours. Next, samples were collected and centrifuged at 1000g for 10 minutes to obtain cell free plasma and conditioned media of PBMCs. Supernatants were passed through a 0.22µm filter before storage at -20°C until further use. To assess the cytokine profile of the different conditions plasma and PBMC conditioned media from 3 donors were pooled separately and subjected to the Proteome Profiler Assay Human XL Cytokine Array Kit (R&D Systems, Minneapolis, MN, USA). The assay was performed according to manufacturer’s instructions. Signals were developed using the Gel Doc XR + device (Bio-Rad Laboratories, Inc., Hercules, California, USA) and dot densities for each cytokine duplicate were calculated using the volume tool in ImageLab 6.0.1 (Bio-Rad Laboratories, Inc.). For visualization, the bar plot option of GraphPad Prism was used (v.8.0.1; GraphPad Software, LA Jolla, CA, USA) to display differences detected in the ATG-treated plasma and PBMC supernatant as fold change relative to the controls. In addition, we measured IFN-γ in supernatants of ATG-treated whole blood, purified PBMCs and purified CD8^+^ T cells as well as human granzyme B in co-cultures via ELISA (Bio-techne, R&D, Minneapolis, Minnesota, USA).

### Flow cytometry

Flow cytometric was performed on a BD FACSCanto II flow cytometer and data were analyzed using FlowJo (10.6.2) software (Tree Star, Ashland, OR). A list of the antibodies used for the detection of cellular epitopes of interest is provided in supplementary table 1.

### INF-γR1 and IFN-γR2 blockade

Isolated monocytes were cultured at 1×10^6^ cell/ml in cell culture medium in the presence of anti-IFN-γR1 antibody (Bio-techne) for 2 hours. Monocytes without added antibody served as controls. Next, conditioned medium obtained from untreated, Isotype-treated and ATG-treated PBMCs was added in the ratio of 1:10. Cells were cultured for a total of 24 hours before flow cytometric assessment of PDL-1 surface expression on monocytes.

### Inhibition of JAK/STAT signaling in purified Monocytes

Purified monocytes were treated with 10µm of the JAK/STAT-Inhibitor ruxolitinib (MedChemExpress, New Jersey, Princeton, USA) diluted in DMSO (Dimethyl sulfoxide, Merck, Darmstadt, germany) for 1 hour before addition of conditioned media of Untreated, Isotype-treated and ATG-treated PBMCs at 1:10. After two hours one replicate of monocytes was lysed in 1x Laemmli Buffer (Bio-Rad Laboratories, Inc.) supplemented with Protease and Phosphatase Inhibitor for assessment of total Stat 1(Cell Signaling Technology, Danvers, Massachusetts, USA) as well as phosphorylated-Stat 1 (Cell Signaling Technology) by western blot. The remaining cells were cultured up to 24 hours before determining surface expression of PDL-1 via flow cytometry.

### Western blotting

Purified Monocytes were lysed in 1x Laemmli Buffer (Bio-Rad Laboratories, Inc.) supplemented with Protease and Phosphatase Inhibitor and loaded on 4–15% SDS-PAGE gels (Bio-Rad Laboratories, Inc.). Proteins were transferred on a nitrocellulose membrane (Bio-Rad Laboratories, Inc.), blocked in non-fat milk with 0.1% Tween20 (Sigma-Aldrich) for 1 hour, and incubated with antibodies as indicated in supplementary table 1 at 4 °C overnight. After washing, membranes were incubated with horseradish-peroxidase conjugated secondary antibodies for 1 hour at room temperature. Signals were developed with SuperSignal West Dura substrate (ThermoFisher Scientific) and imaged with a Gel Doc XR + device (Bio-Rad Laboratories, Inc.). Quantification analysis was performed using the Volume tool in ImageLab 6.0.1 (Bio-Rad Laboratories, Inc.).

### Co-culture of activated CD8+ T cells with monocytes

CD8^+^ T cells were isolated from whole blood using CD8^+^ magnetic beads (Miltenyi) and the quadroMACS separator (Miltneyi). 5×10^6^ cells/ml were stained with CellTrace^™^ violet proliferation kit (ThermoFisher Scientific) according to manufacturer’s instructions. Next, anti-CD3 and anti-CD28 T cell activation beads were added to the isolated T cells before placing them in a round-bottom 96 well plate. A total of 5×10^5^ cells was pipetted into each well. After 24 hours of activation the same amount of autologous monocytes that had meanwhile been treated with conditioned media of untreated PBMCs and ATG-treated PBMCs were added to the T lymphocytes. In addition, we incubated PDL-1^+^ monocytes with for 1 hour with durvalumab (Imfinzi, Medimmune, Astra Zeneca, Cambridge, UK), a monoclonal antibody directed against PDL-1, to assess its contribution on inhibition of CD8^+^ T cell proliferation. Experiments were performed with 4 independent donors. After 4 days of co-culture cells were collected stained for CD8-PECy5 (BioLegend) and assessed for cell proliferation with flow cytometry. Conditioned media of co-cultured cells were collected and stored at -20°C until further use.

### Statistical analysis

For single cell RNA seq, two donors were analyzed. Negative binomial regression was performed to normalize data and achieve variance stabilization. Wilcoxon rank sum test was followed by Benjamini-Hochberg post-hoc test to calculate differentially expressed genes. For *in vitro* experiments, at least three different donors were used. Data were statistically evaluated using GraphPad Prism v8.0.1 software (GraphPad Software, San Diego, USA). When analyzing three or more groups ordinary one-way ANOVA and multiple comparison post hoc tests with Dunnett’s correction were calculated. with p-values < 0.05 considered as statistically significant. Data are presented as mean ± standard error of the mean (SEM).

## Supplementary Materials

Fig. S1. Comparison of differential responses of cytotoxic *CD8*^*+*^ T cells and myeloid cells

Fig. S2. Differential gene expressions of immune cells from human whole blood treated with ATG

Fig. S3. Cytokine profiling of PBMCs treated with ATG.

Fig. S4. Gating strategies for flow cytometry experiment.

Fig. S5. Increased surface expression of PDL-1 on monocytes monitored over time.

Fig. S6. Leading enrichment genes for regulation of IFN-γ mediated signaling pathway in monocytes treated with ATG

Fig. S7. Blockage of IFN-γR2 does not prevent induction of PDL-1 on monocytes stimulated with conditioned medium of ATG treated PBMCs.

Fig. S8. Differential regulation of JAK/STAT pathway members across lymphoid and myeloid immune cells after treatment with ATG

Fig. S9. Pretreatment of PBMCs with hydrocortisone does not prevent induction of PDL-1 by ATG.

Table S1. Overview of used antibodies

## Acknowledgments

We would like to thank Dr. Hans Peter Haselsteiner and the CRISCAR Familienstiftung for their ongoing support of the Medical University of Vienna/ Aposcience AG public private partnership aiming to augment basic and translational clinical research in Austria/Europe. The authors acknowledge the core facilities of the Medical University of Vienna, a member of Vienna Life Science.

## Author contributions

Conceptualization: DC, HJA, MM; Methodology: DC, KK, ML, MD, DB; Investigation: DC, MM; Visualization: DC, DB; Funding acquisition: HJA, MM; Project administration: HJA, MM; Supervision: HJA, MM; Writing – original draft: DC, MM; Writing – review & editing: KK, MD, DB, ML, HJA

## Data and materials availability

Except for scRNA-seq experiment all data are available in the main text or the supplementary materials. The scRNA-seq data are accessible upon request

