## Supplementary figures for "Antithymocyte globulin inhibits CD8^+^ T cell effector functions via the paracrine induction of PDL-1 on monocytes"

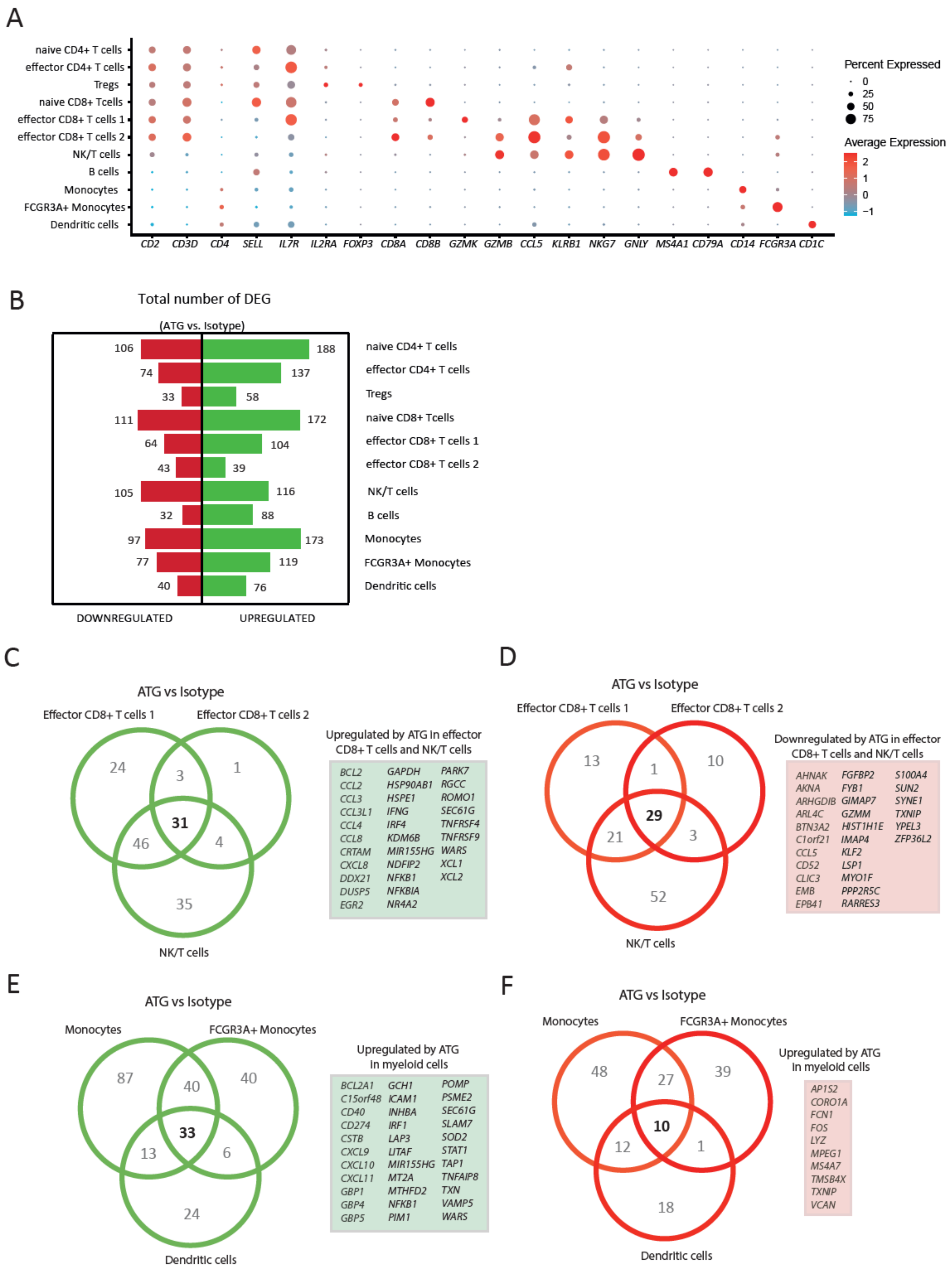
A
naive CD4+ T cells
effector CD4+ T cells
Tregs
naive CD8+ Tcells
effector CD8+ T cells 1
effector CD8+ T cells 2
NK/T cells
B cells
Monocytes
FCGR3A+ Monocytes
Dendritic cells
CD2 CD3D CD4 SELL IL7R IL2RA FOXP3 CD8A CD8B GZMK GZMB CCL5 KLRB1 NKG7 GNLY MS4A1 CD79A CD14 FCGR3A CD1C
Percent Expressed
0
25
50
75
Average Expression
2
1
0
−1
B
Total number of DEG
(ATG vs. Isotype)
106 188 naive CD4+ T cells
74 137 effector CD4+ T cells
33 58 Tregs
111 172 naive CD8+ Tcells
64 104 effector CD8+ T cells 1
43 39 effector CD8+ T cells 2
105 116 NK/T cells
32 88 B cells
97 173 Monocytes
77 119 FCGR3A+ Monocytes
40 76 Dendritic cells
DOWNREGULATED
UPREGULATED
C
ATG vs Isotype
Effector CD8+ T cells 1
Effector CD8+ T cells 2
24 3 1 31 46 4 35
NK/T cells
Upregulated by ATG in effector CD8+ T cells and NK/T cells
BCL2 CCL2 CCL3 CCL3L1 CCL4 CCL8 CRTAM CXCL8 DDX21 DUSP5 EGR2
GAPDH HSP90AB1 HSPE1 IFNG IRF4 KDM6B MIR155HG NDFIP2 NFKB1 NFKBIA NR4A2
PARK7 RGCC ROMO1 SEC61G TNFRSF4 TNFRSF9 WARS XCL1 XCL2
D
ATG vs Isotype
Effector CD8+ T cells 1
Effector CD8+ T cells 2
13 1 10 29 21 3 52
NK/T cells
Downregulated by ATG in effector CD8+ T cells and NK/T cells
AHNAK AKNA ARHGDIB ARL4C BTN3A2 C1orf21 CCL5 CD52 CLIC3 EMB EPB41
FGFBP2 FYB1 GIMAP7 GZMM HIST1H1E IMAP4 KLF2 LSP1 MYO1F PPP2R5C RARRES3
S100A4 SUN2 SYNE1 TXNIP YPEL3 ZFP36L2
E
ATG vs Isotype
Monocytes
FCGR3A+ Monocytes
87 40 40 33 13 6 24
Dendritic cells
Upregulated by ATG in myeloid cells
BCL2A1 C15orf48 CD40 CD274 CSTB CXCL9 CXCL10 CXCL11 GBP1 GBP4 GBP5
GCH1 ICAM1 INHBA IRF1 LAP3 LITAF MIR155HG MT2A MTHFD2 NFKB1 PIM1
POMP PSME2 SEC61G SLAM7 SOD2 STAT1 TAP1 TNFAIP8 TXN VAMP5 WARS
F
ATG vs Isotype
Monocytes
FCGR3A+ Monocytes
48 27 39 10 12 1 18
Dendritic cells
Upregulated by ATG in myeloid cells
AP1S2 CORO1A FCN1 FOS LYZ MPEG1 MS4A7 TMSB4X TXNIP VCAN

**Fig. S1. Comparison of differential responses of cytotoxic *CD8*$^+$ T cells and myeloid cells to ATG**

**(A)** Dotplot with cluster marker genes for cluster of differentiation (*CD*) genes *CD2*, *CD3D*, *CD4*, *CD8A*, *CD8B*, *CD79A*, *CD14*, *CD1C* and functional state defining genes including Selectin L (*SELL*), Interleukin 7 receptor (*IL7R*), Interleukin 2 receptor subunit alpha (*IL2RA*), Forkhead Box P3 (*FOXP3*), Granzyme K (*GZMK*), Granzyme B (*GZMB*), C-C Motif Chemokine Ligand 5 (*CCL5*), Killer Cell Lectin Like Receptor B1 (*KLRB1*), Natural Killer Cell Granule Protein 7 (NKG7), Granulysin (*GNLY*), Membrane Spanning 4-Domains A1 (*MS4A1*) and Fc Gamma Receptor IIIa (*FCGR3A*). Percentage of cells expressing the respective gene is illustrated by dot size while color gradient from light blue (low) to red (high) highlights average expression. **(B)** Bar graph with absolute numbers of differentially up- (green) and downregulated (red) genes for all identified cell types. Venn diagrams displaying shared (C) up- and (D) downregulated genes for **(C & D)** effector *CD8*$^+$ T cells 1, effector *CD8*$^+$ T cells 2 and NK/T cells as well as **(E & F)** monocytes, FCGR3A+ monocytes and dendritic cells. Respective genes are provided in boxes.

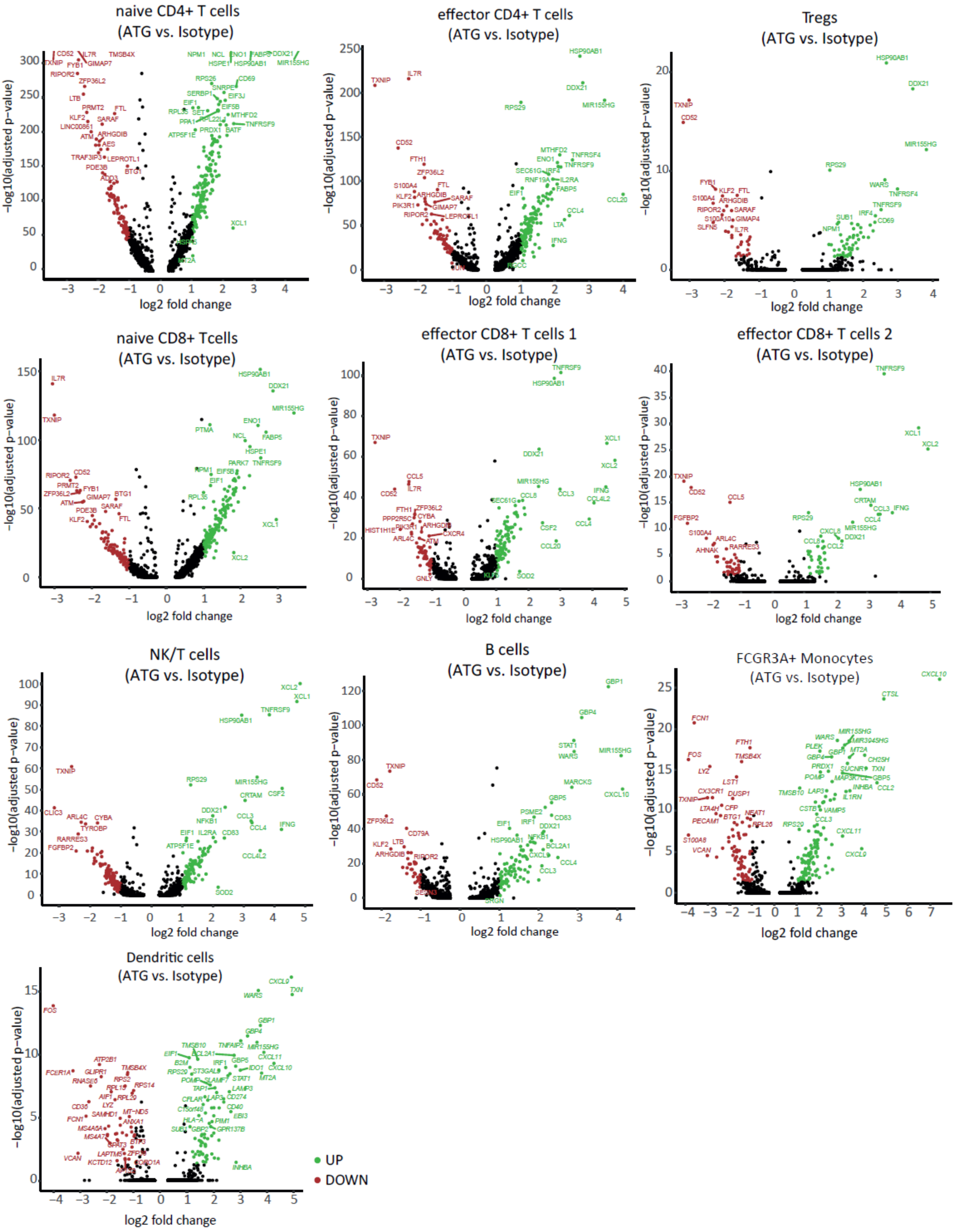
naive CD4+ T cells
(ATG vs. Isotype)
effector CD4+ T cells
(ATG vs. Isotype)
Tregs
(ATG vs. Isotype)
naive CD8+ Tcells
(ATG vs. Isotype)
effector CD8+ T cells 1
(ATG vs. Isotype)
effector CD8+ T cells 2
(ATG vs. Isotype)
NK/T cells
(ATG vs. Isotype)
B cells
(ATG vs. Isotype)
FCGR3A+ Monocytes
(ATG vs. Isotype)
Dendritic cells
(ATG vs. Isotype)
−log10(adjusted p−value)
log2 fold change
UP
DOWN

**Fig. S2. Differentially expressed genes across lymphoid and myeloid cell types after treatment with ATG.**

Volcano plots depicting differentially up- (green) and downregulated (red) genes with annotations.

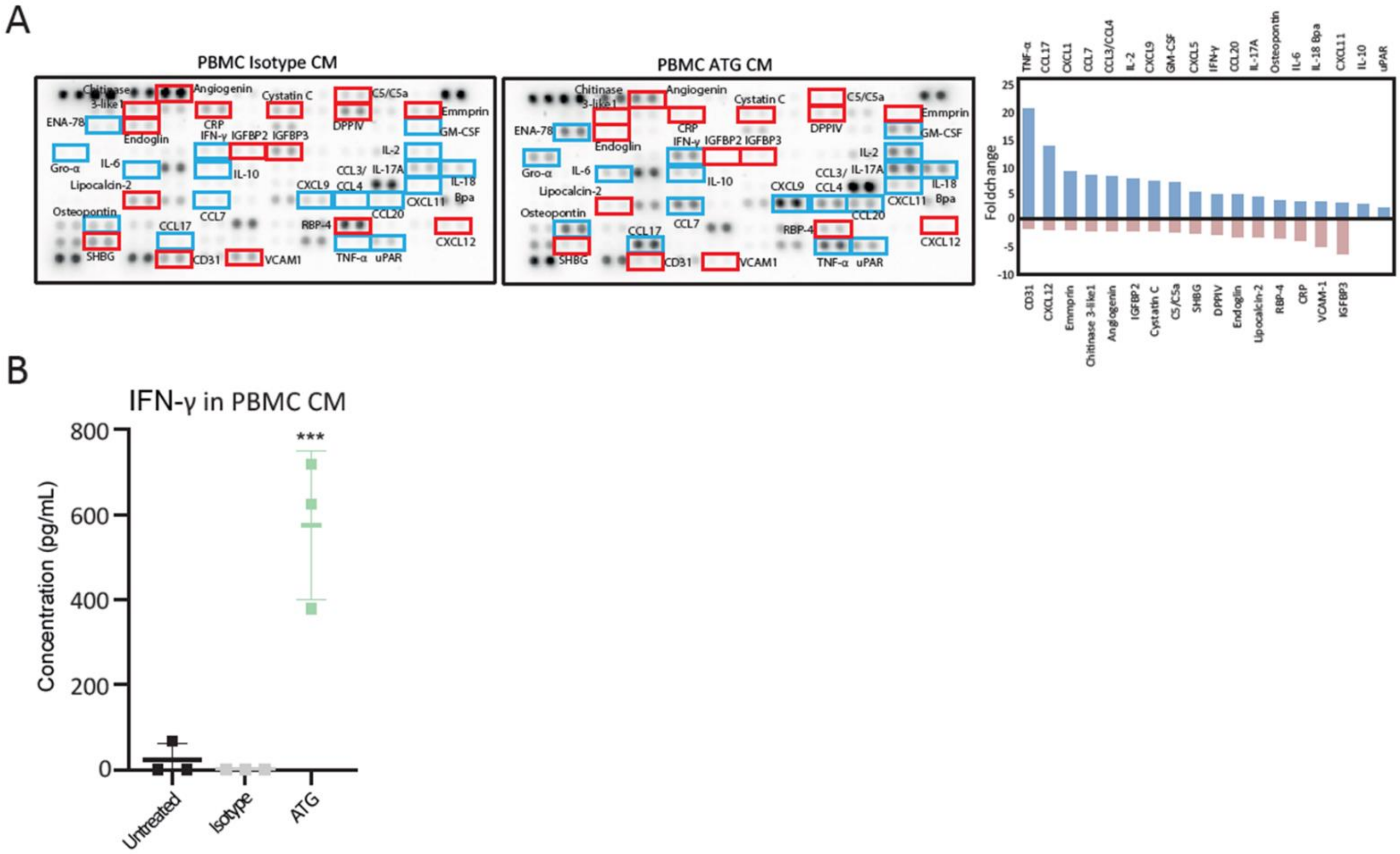


**Fig. S3. Cytokine profiling of PBMCs treated with ATG.**

**(A)** Immunodetection array membrane of conditioned medium of purified PBMCs treated with ATG and isotype control. Pooled supernatants of 3 donors per condition were analyzed. Proteins with a > 2-fold increase are shown in blue brackets while proteins with a fold change < 0.5 are marked in red brackets. Bar plots highlight the differentially detected proteins. Each colored bar represents one analyte. **(B)** Concentration of IFN-γ in conditioned media of isolated PBMCs. N = 3 donors, ANOVA was used to determine statistically significant differences in IFN-γ concentrations between conditions with a p-value = 0.0008.

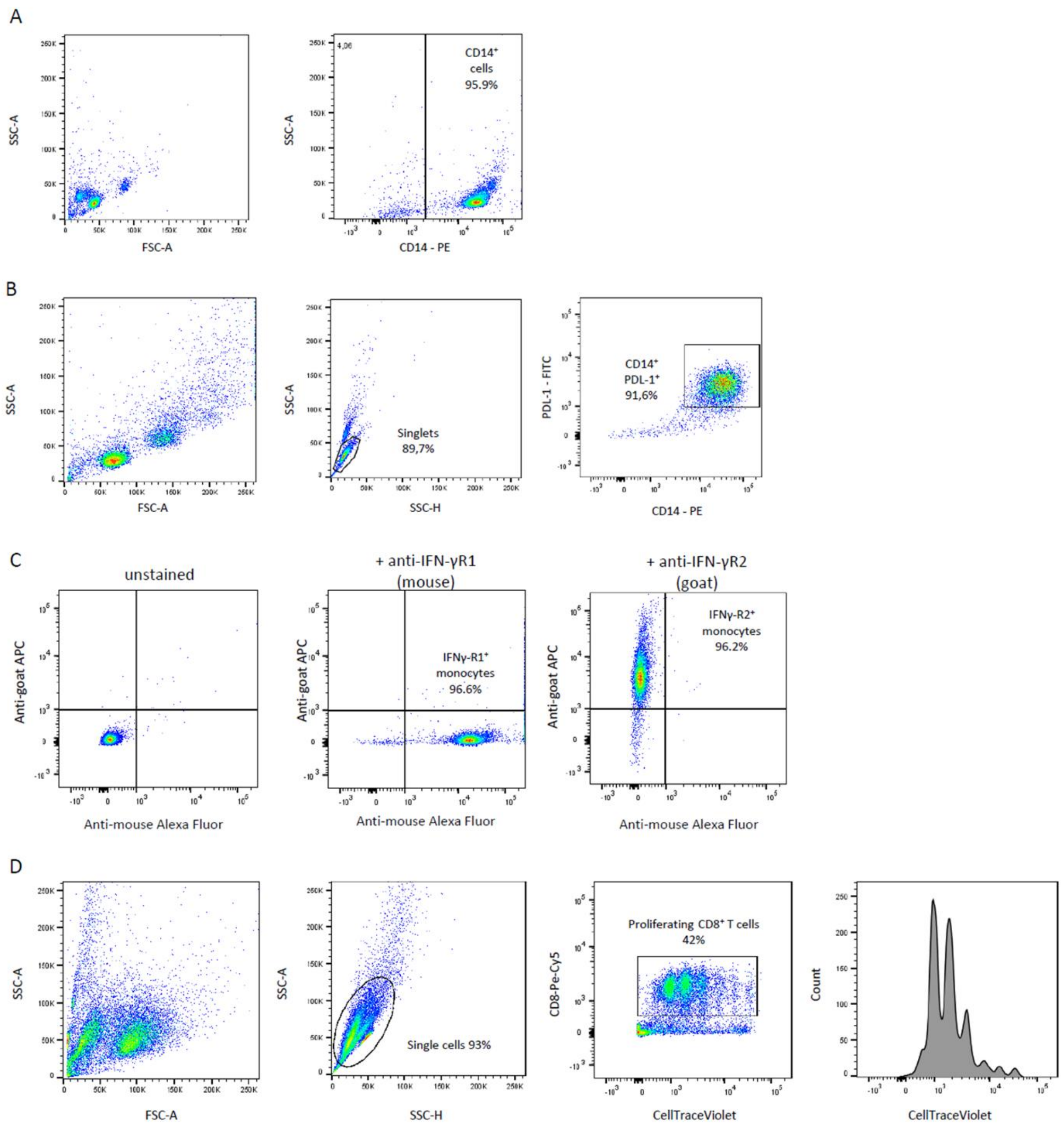


**Fig. S4. Gating strategy for flow cytometry experiments.**

Gating strategies for **(A)** purity of monocytes, **(B)** PDL-1 status of purified monocytes, **(C)** confirmation of IFNγ-R1 surface expression on monocytes and **(D)** co-culture of purified CD8+ T cells and monocytes.

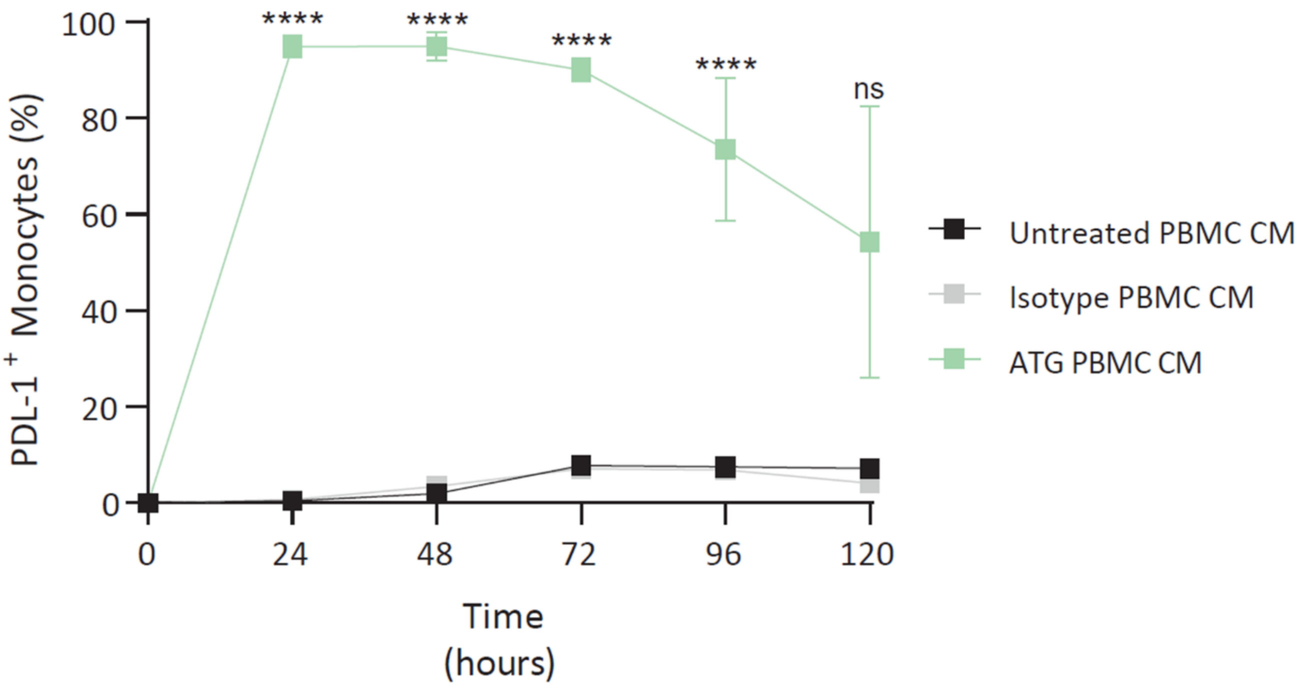


**Fig. S5. Increased surface expression of PDL-1 on monocytes monitored over time.** Monocytes were treated with conditioned medium of ATG stimulated PBMCs and assessed for percentage PDL-$1^{+}$- monocytes every 24 hours on 5 consecutive days. Expression of PDL-1 was significantly higher up to 96 hours after a single administration of the conditioned medium. For the time points of 24h, 48h and 72h we registered a p-value < 0.0001 while at 96h p-value = 0.0081. No statistically significant differences in the mean percentage of PDL-1+ monocytes were detected between conditions after 120h. N = 3

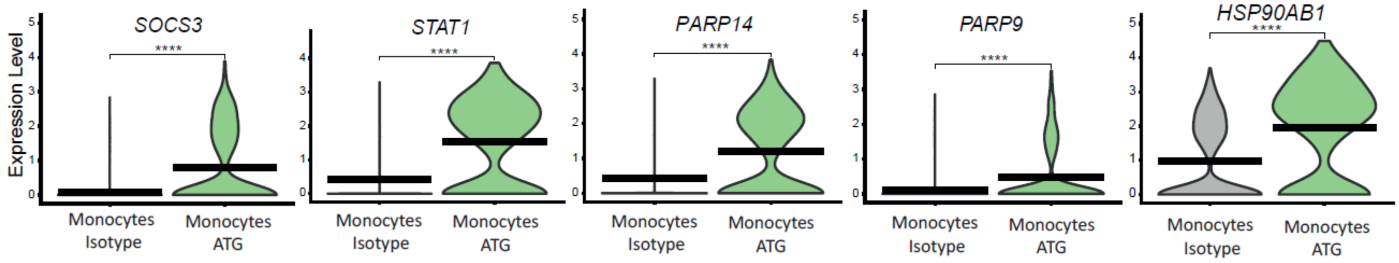


**Fig. S6. Leading enrichment genes for regulation of IFN-γ mediated signaling pathway in monocytes treated with ATG.**

Violin plots depicting core enrichment genes differentially expressed between monocytes Isotype and monocytes ATG for *SOCS3*, *STAT1*, *PARP14*, *PARP9*, *HSP90AB1*. Expression levels are indicated by violin plot height while width represents proportion of positive cells. Crossbars mark mean expression. **** indicate p-value < 0.0001

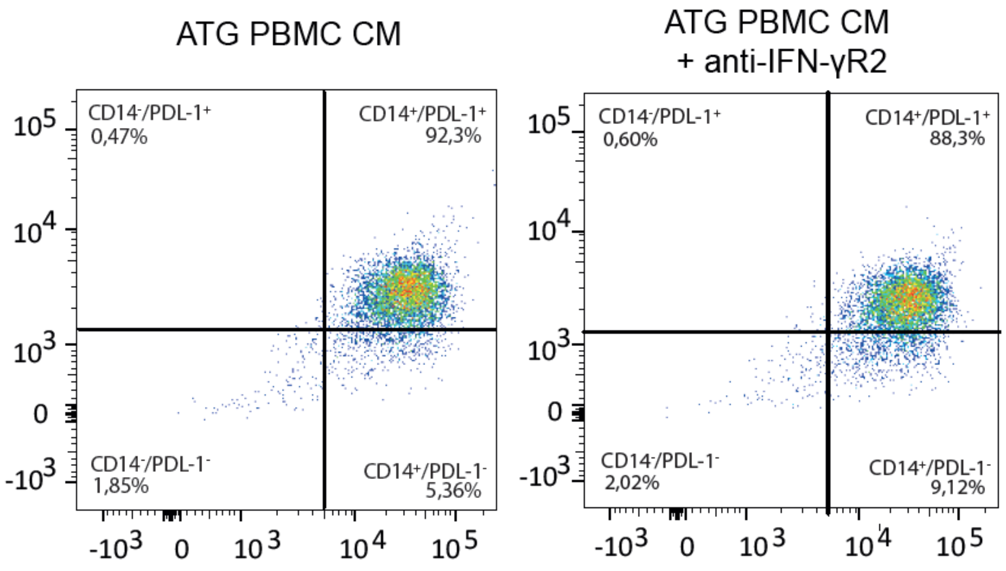


**Fig. S7. Blockage of IFN-γR2 does not prevent induction of PDL-1 on monocytes stimulated with conditioned medium of ATG treated PBMCs.**

FACS plots of monocytes with and without pre-incubation with antibodies to IFN-γR2 before treatment with conditioned medium.

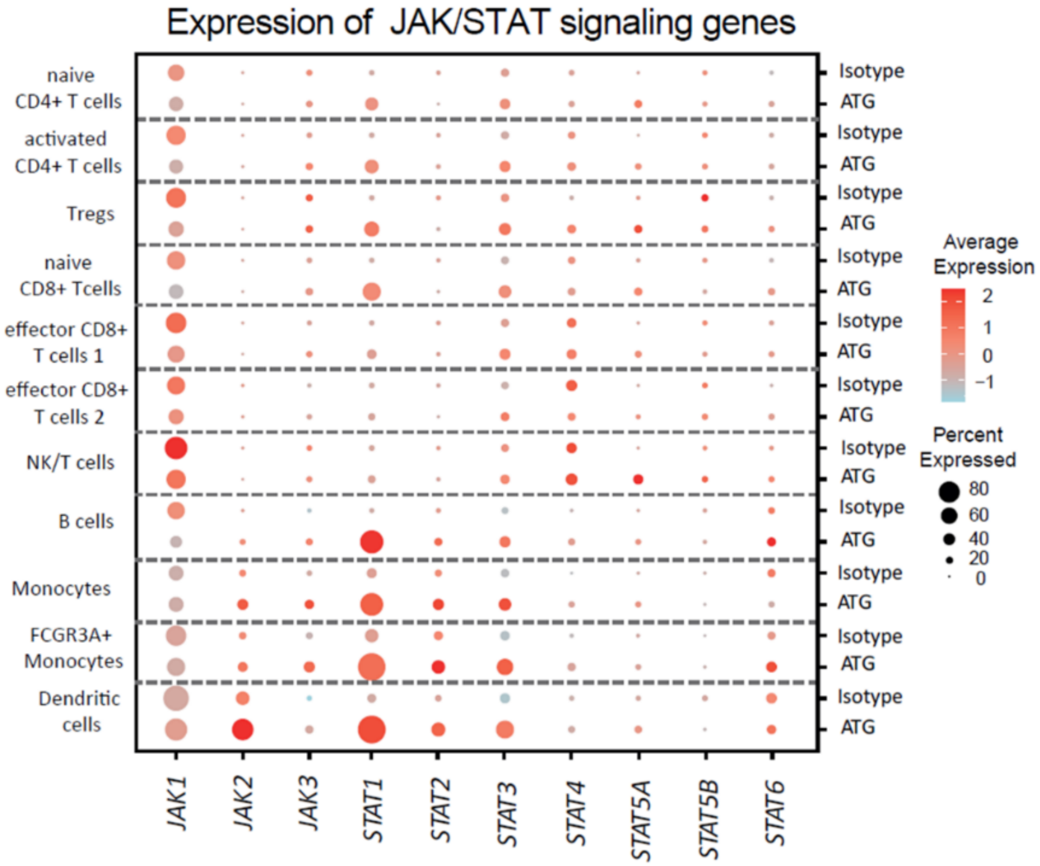


**Fig. S8. Differential regulation of JAK/STAT pathway members across lymphoid and myeloid immune cells after treatment with ATG**

Overview on JAK/STAT genes and their regulations by ATG compared to Isotype across all cell types. Dot size illustrates percentage of cells expressing the respective gene while color gradient from light blue (low) to red (high) highlights average expression.

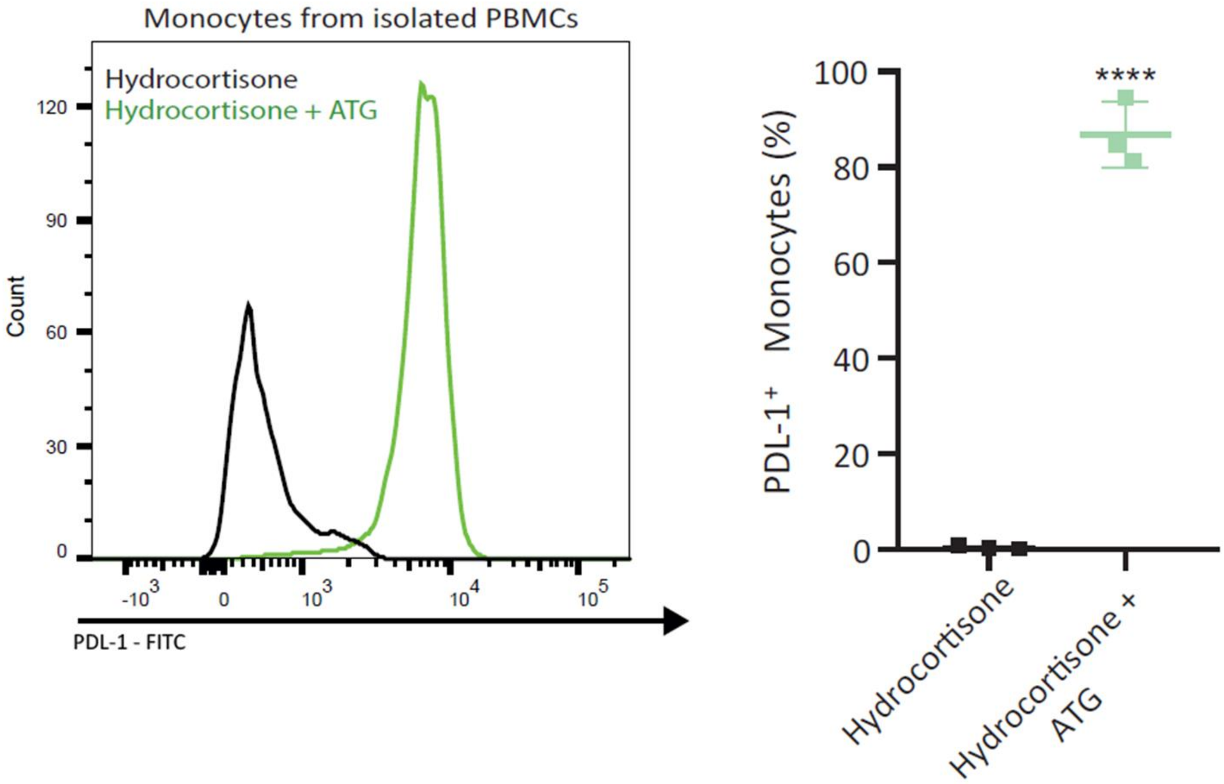


**Fig. S9. Pretreatment of PBMCs with hydrocortisone does not prevent induction of PDL-1 by ATG.**

Histogram displaying PDL-1+ monocytes from PBMCs treated with 10 mM hydrocortisone for 2 hours prior to addition of ATG. After 24 hours monocytes were assessed for PDL-1. Percentage of PDL-1+ monocytes was significantly lower in monocytes from PBMCs treated only with hydrocortisone (0.37 ± 0.19%) when compared to the combination of hydrocortisone and ATG (86.8 ± 4%, p-value < 0.0001). N = 3

**Table 1. Overview on used antibodies.**

| Target | Species | Conjugate | Catalog #/ RRID | Company | Dilution/Concentration | Comment |
|---|---|---|---|---|---|---|
| **Flow cytometry** | | | | | | |
| CD14 | Rabbit | PE | 325605 | BioLegend | 1:50 | 20 min, RT |
| CD8 | Rabbit | PE-Cy5 | 344770 | BioLegend | 1:50 | 20 min, RT |
| PDL-1 | Rabbit | FITC | 393606 | BioLegend | 1:50 | 20 min, RT |
| **Western blot** | | | | | | |
| Stat 1 | Rabbit | n.a. | 9172 | Cell Signaling | 1:200 | o.n., 4°C |
| p-Stat 1 (Y701) | Rabbit | n.a. | 9167S | Cell Signaling | 1:200 | o.n., 4°C |
| **Blocking experiments** | | | | | | |
| IFN-γR1 | Mouse | n.a. | MAB6731 | Bio-Techne | 1:100 | 2 hr, 37°C |
| IFN-γR2 | Goat | n.a. | AF773SP | R&D Systems | 2.5 μg/ml | 2 hr, 37°C |
| **Secondary antibodies** | | | | | | |
| Anti-rabbit | Goat | HRP | 1706515 | Bio-Rad | 1:10000 | 1 hr, RT |
| Anti-mouse | Goat | Alexa Fluor | A-21151 | ThermoFisher Scientific | 1:250 | 20 min, RT |
| **T cell activation** | | | | | | |
| NA/LE Anti-human CD3<br><br>Clone OKT3 | Mouse | n.a. | AB_2869821 | BD Biosciences | 3 μg/ml | 24 hr, 37°C |
| NA/LE Anti-human CD28<br><br>Clone CD28.2 | Mouse | n.a. | AB_396068 | BD Biosciences | 5 μg/ml | 24 hr, 37°C |
